# Decisions are expedited through multiple neural adjustments spanning the sensorimotor hierarchy

**DOI:** 10.1101/203141

**Authors:** Natalie A. Steinemann, Redmond G. O’Connell, Simon P. Kelly

**Affiliations:** Department of Biomedical Engineering, The City College of The City University of New York, New York, NY, 10031; Zuckerman Mind Brain Behavior Institute, Columbia University, New York, NY 10023; Trinity College Institute of Neuroscience and School of Psychology, Trinity College Dublin, Dublin 2, Ireland; School of Electrical and Electronic Engineering, University College Dublin, Dublin 4, Ireland

## Abstract

When decision makers prioritize speed over accuracy, neural activity is elevated in brain circuits involved in preparing actions. Such “urgency” signal components, defined by their independence from sensory evidence, are observed even before evidence is presented and can grow dynamically during decision formation. Is urgency applied globally, or are there adjustments of a distinct nature applied at different processing levels? Using a novel multi-level recording paradigm, we show that dynamic urgency impacting cortical action-preparation signals is echoed downstream in electromyographic indices of muscle activation, but does not directly influence upstream cortical levels. A motor-independent representation of cumulative evidence reached lower pre-response levels under conditions of greater motor-level urgency, paralleling a decline in choice accuracy. At the sensory level itself, we find a boost in differential evidence, which is correlated with changes in pupil size and acts to alleviate, rather than contribute to, the overall accuracy cost under speed pressure.

---

When situations call for it, animals can prioritize speed over accuracy in their sensory-guided actions. Prominent computational models suggest that sensorimotor decisions are made by drawing sequential samples from noisy evidence representations and integrating them up to an action-triggering threshold^1,2^. In this framework speed can be emphasized at the expense of accuracy by lowering this threshold, which in the models may be constant or collapsing (i.e., narrowing) over the timeframe of the decision^3,4^. Neural circuits involved in preparing decision-reporting actions have been found to implement such adjustments in the form of “urgency” signal components, which non-selectively elevate activity towards action thresholds. A “static” component of urgency has been widely observed in raised baseline activity before evidence presentation^5–8^, and recent work has further revealed a “dynamic” component that grows over the course of a decision, effectively implementing a collapsing bound^8–11^. A key defining property of urgency is that it is generated purely from knowledge of time constraints and/or elapsed time itself, and it contributes to neural buildup activity alongside, but strictly independent of, the influence of sensory evidence^12^. This means that any speed benefits of urgency necessarily incur a cost to choice accuracy. Thus far, urgency components have been identified only in neural circuits involved in preparing actions. Recent work has implicated diffusely-projecting neuromodulatory systems in the generation of urgency^11,13^, suggesting that it may, in fact, act globally, i.e., at all levels of the sensorimotor hierarchy. However, this remains untested. Are speed pressure adjustments applied at processing levels other than effector-selective action preparation, and if so, do they act as evidence-independent urgency components, to the detriment of accuracy?

The impact of speed pressure at the level of sensory processing is particularly unclear. Theoretical work has suggested that increased cognitive “effort” during time-pressured decisions could enhance the efficiency^14^ or reliability^15^ of sensory evidence encoding. However, in the rare instances where computational model fits have indicated that speed pressure affects the evidence strength-dependent parameter of drift rate, it has been decreased, suggesting a weakening of sensory representations^16,17^. On the other hand, in monkeys performing visual search, salience-encoding visual neurons of the frontal eye field (FEF), have exhibited increased and earlier target selectivity under speed pressure, consistent with a stronger evidence representation in this scenario^18^. Yet a different picture is painted by human fMRI studies, which have indicated an unchanged^5^ or decreased^19^ influence of sensory evidence on decision formation under speed pressure and report no changes in activation at the sensory level itself.

Recent studies have highlighted the existence of abstract, motor-independent processes that intermediate between sensory evidence encoding and motor preparation, and afford flexibility in the mapping of one to the other^20–22^, and such abstract properties have been demonstrated for dynamic-evidence accumulation signatures in humans^23^. The impact of speed pressure on such signals is unknown, yet it stands to be highly illuminating on the nature and utility of effector-general decision signals in the brain. On one hand, if not at the sensory level itself, it is conceivable that urgency could be applied *de novo* at this abstract stage of integration and inherited by downstream effector-specific signals. This would constitute an artificial form of "evidence" in itself, hastening the process of cognitive deliberation and not just the preparation of actions. Alternatively, urgency signals may first confluence with evidence at downstream motor levels, thus allowing an unadulterated representation of cumulative evidence to be retained at the motor-independent level.

Resolving all of these questions requires a global, system-wide view over all levels of processing in the multilayered neural architecture for decision-making^24–28^. Here, we employed a novel contrast-comparison decision paradigm that enables neural dynamics at the key hierarchical processing levels to be traced simultaneously in humans making decisions under varying response time constraints. Using scalp electroencephalography (EEG), we traced sensory evidence encoding via stimulus-driven steady-state visual evoked potentials (SSVEP) reflecting contrast-dependent responses in early visual cortex^29^. We traced motor preparation in effector-selective spectral amplitude changes in the Mu/Beta band (8-30 Hz) over the motor cortex contralateral to prepared thumb-press responses^30^. Like signals previously identified in sensorimotor neurons of monkey and rodent brains^1,31^, Mu/Beta reflects the key characteristics of a theoretical decision variable, namely a build-up rate that scales with the strength of sensory evidence (e.g. dot motion coherence) and a bound-crossing relationship to response execution^23,32,33^. Also like sensorimotor decision signals in monkey^8^, these spectral EEG measures of motor preparation have been shown to reflect both static and dynamic components of evidence-independent urgency^11^. To determine whether effects of speed pressure at this cortical level translate downstream to the peripheral level of muscle activation, we recorded electromyographic (EMG) signals from both alternative response effectors (left/right thumbs). To measure upstream, motor-independent representations of cumulative evidence, we traced the timecourse of a recently characterized centro-parietal positivity (CPP) in the event-related potential (ERP). Like motor preparation signals, the CPP has been shown to build at a rate that scales with evidence strength and to peak around the time of the response, regardless of the sensory feature or modality being decided upon^23,33^. However, several important distinctions have established its more abstract, motor-independent nature. When detection decisions were covertly counted rather than immediately reported, the CPP continued to trace evidence accumulation despite the absence of motor responses, while Mu/Beta changes were absent, verifying that there was no motor preparation^23^. In a task requiring delayed decision reports, the CPP rose and fell during evidence presentation consistent with early completion of the sensory decision itself, whereas effector-selective motor preparation was sustained until the decision-reporting action was eventually permitted^34^. Moreover, when the stimulus-response mapping was unknown during evidence presentation and only revealed later, the CPP exhibited the same rise and fall despite selective motor preparation being wholly absent in this case^34^. Based on these features, along with the fact that the CPP buildup temporally precedes that of evidence-selective motor preparation^33^, it has been suggested that the CPP lies at an intermediate, motor-independent level in the decision hierarchy, which receives sensory evidence as input and in turn feeds into motor preparation.

We found multiple distinct adjustments for speed pressure across the hierarchy. Cortical motor preparation signals exhibited static and dynamic urgency signal components. Downstream, peripheral muscle activation was shortened and intensified, and reflected dynamic urgency in an increase in evidence-independent activation over the course of a decision. Upstream, we found a qualitatively distinct speed-pressure modulation at the sensory level that rendered the alternatives more discriminable but had no evidence-independent component. This differential boost was correlated with neuromodulatory influences reflected in pupil size and was linked to improved response accuracy. The motor-independent CPP was spared from a direct influence of evidence-independent urgency, but was curtailed in the level it reached at response by the downstream, motor-level urgency. Thus, speed pressure induces adjustments at multiple levels of the sensorimotor hierarchy, but of qualitatively distinct nature; while urgency applied at the latter motor-related levels expedites responses at the expense of accuracy, adjustments at the sensory front-end work to alleviate some of this accuracy cost, and allow an unadulterated rendering of the cumulative, bottom-up evidence at the intermediate, motor-independent level.

## Results

Sixteen human participants performed a two-alternative forced-choice contrast discrimination task at two different, interleaved evidence strengths. Subjects were instructed to report whether the left- or right-tilted lines in a compound overlay-pattern had a greater contrast by pressing a button with the thumb of the corresponding hand (Figure 1a). On a trial-by-trial basis, participants were incentivized to emphasize decision speed or accuracy, as indicated by a color cue at the beginning of the trial (see Online Methods and Supplementary Fig. 1). Randomized trial-by-trial cueing was used so that the short-term establishment of pre-decision states in preparation for speed pressure could be examined at each neural processing level.

### Response accuracy decreases with speed pressure and with reaction time

Responses were faster and more accurate for larger contrast differences (RT: F(1,15)=86.7, p=1.27*10^−7^; accuracy: F(1,15)=106.8, p=3.23*10^−8^ Supplementary Table 2a-b). Subjects made faster responses (F(1,15)=46.6, p=5.71*10^−6^) at the expense of response accuracy (F(1,15)=23.2, p=0.00023) in the Speed regime compared to the Accuracy regime (Figure 1b). Plotting decision accuracy as a function of RT quantiles additionally revealed that extremely fast responses in the Speed regime were especially inaccurate, and that beyond approximately 600ms, accuracy decreased monotonically with increasing RT at a similar rate in both regimes (Figure 1c). To further characterize the accuracy decline over longer RTs, we computed a linear mixed-effects regression model restricted to RT bins lying beyond each subject’s point of maximum response accuracy. Although response accuracy was significantly lower under speed pressure (t(508)=7.6, p=1.34*10^−13^, Supplementary Table 1a), for low contrast stimuli (t(508)=13.6, p=2.63*10^−36^), and for slower response times (t(508)=−15.0; p=2.46*10^−42^), adding a term capturing the interaction between RT and Speed/Accuracy regime did not improve the fit significantly (delta log likelihood=1.8236; p=0.24). Thus, response accuracy declined over RT in the Accuracy as much as the Speed regime.

**FIGURE 1:**
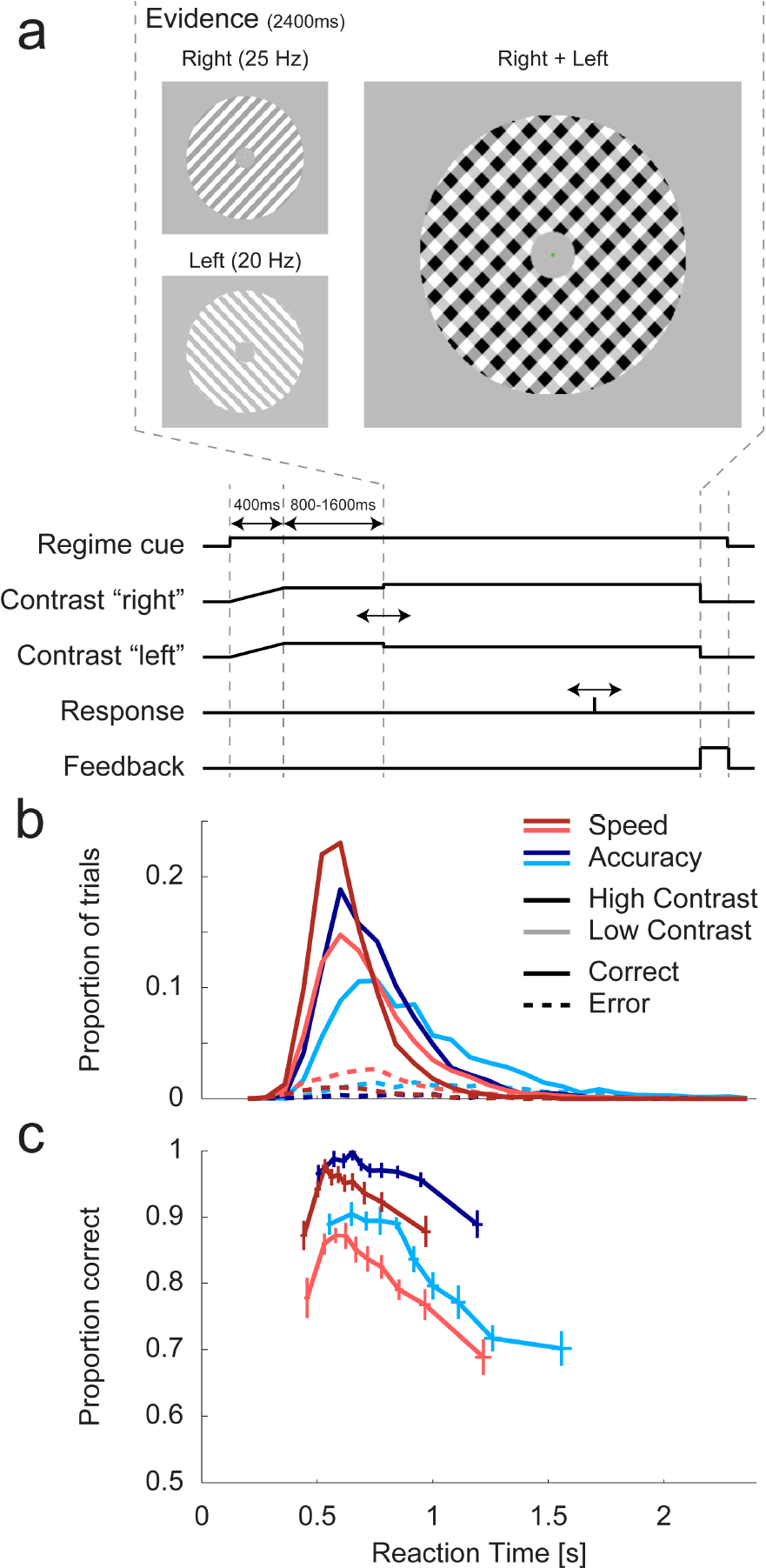
Task and Behavior - RT histograms and conditional accuracy functions. (a) Task structure and trial timing. Two overlaid grating patterns tilted 45 degrees to the left and right were phase-reversed at 20Hz and 25Hz, respectively. Both gratings initially had an equal contrast of 50% (“Baseline”), and after a variable delay one stepped to 56% (low contrast) or 62% (high contrast), while the other decreased by the same amount (44% or 38%). In the example stimulus in panel A the right-tilted pattern increased in contrast. This contrast-difference “evidence” was displayed for 2.4s and evidence onset was marked by an auditory cue to avoid temporal ambiguity. (b) Reaction time distribution for correct (solid) and incorrect (dashed) response trials of high (darker lines) and low (lighter lines) contrast differences in the Speed (red) and Accuracy (blue) regime. (**c**) Response accuracy computed in each of ten reaction time bins separately for each condition. Horizontal and vertical error bars denote the standard error of the mean across subjects. Apart from a low response accuracy for very fast responses, conditional accuracy functions of all conditions declined over RT.

### Enhancement of differential sensory evidence under speed pressure

The neural representation of sensory evidence was quantified as the difference in spectral amplitude (occipital SSVEP) between the flicker frequencies of the two phase-reversing gratings (left-tilted minus right-tilted). As expected, these differential SSVEP signals underwent a change in opposing directions for trials in which the left-versus right-tilted grating was higher in contrast (Figure 2b, thin vs. thick lines; F(1,15)=27.1, p=0.0027, Supplementary Table 2c), and this directional amplitude change was strongly modulated by the difference in stimulus Contrast (F(1,15)=38.4, p=0.000017). More surprisingly, differential evidence was also significantly boosted under speed pressure (F(1,15)=8.8, p=0.0096). This modulation was transient, emerging following evidence onset and lasting until just before the response (Supplementary Table 2d-e), suggesting that this differential boost was invoked specifically during decision formation. SSVEP amplitudes for individual phase-reversal frequencies showed no modulation at baseline (t-tests; 20Hz: t(15)=0.43, p=0.68; 25Hz: t(15)=0.52, p=0.61) or main effect of speed pressure during decision formation (p>0.1 for each individual frequency in the time range of significant Speed/Accuracy x Left/Right interaction; See Supplementary Fig. 2 for time-resolved analysis, Supplementary Table 2f-g), indicating that this differential evidence boost did not arise from a general, multiplicative boost in amplitude to both sensory components, but, like the task itself, was truly differential in nature. Further, correct trials were associated with greater differential SSVEP amplitude (higher-contrast grating minus lower-contrast) than error trials even accounting for the factors of Contrast, Target type and Speed/Accuracy regime, indicating that the impact of boosting the differential evidence signal is to improve accuracy as well as speed (linear mixed-effects model; t(15109) = 2.57; p = 0.010153, Supplementary Table 1b).

**FIGURE 2:**
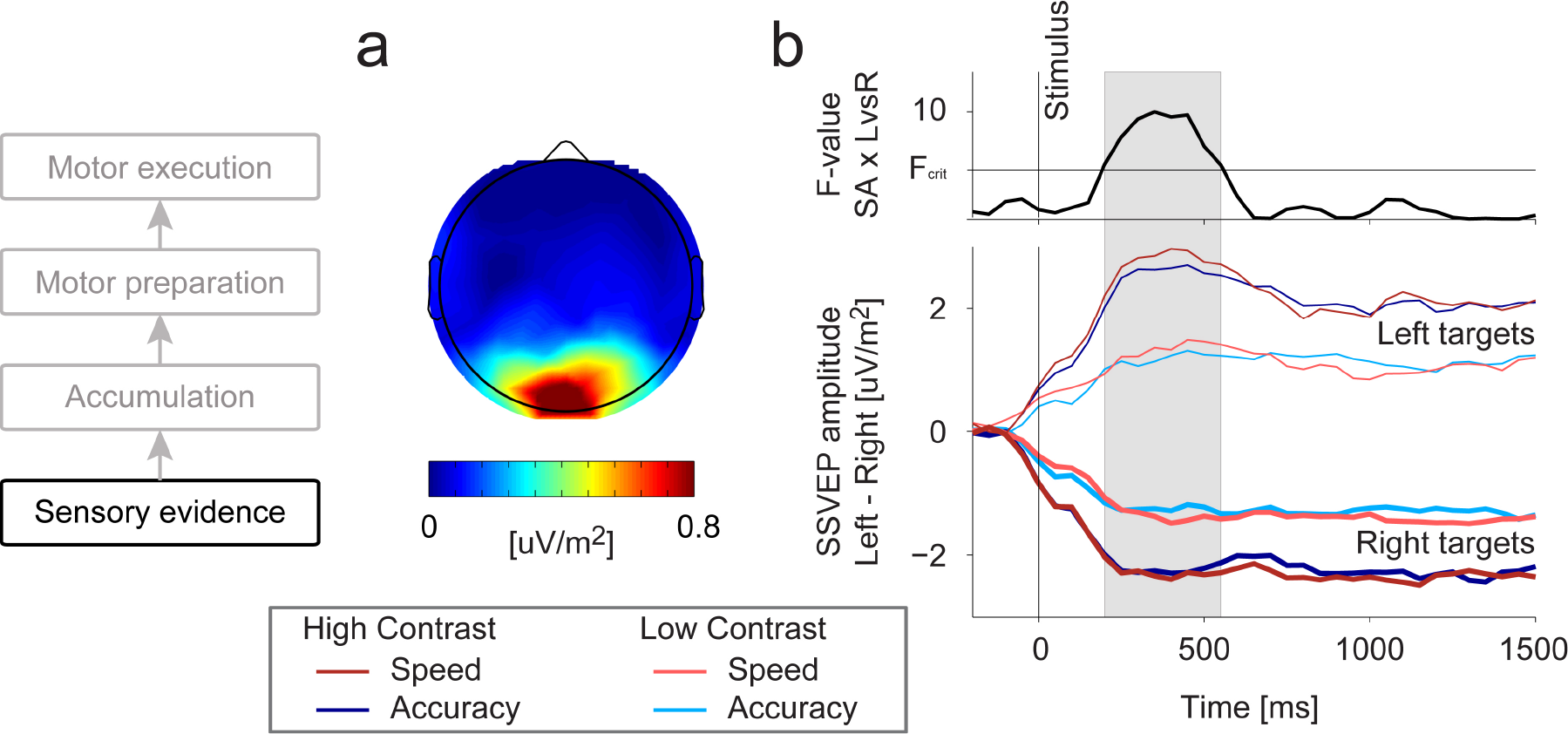
Sensory evidence modulations due to speed pressure. (**b**) Average steady-state visual evoked potential (SSVEP) measured in the 800ms before stimulus-onset was maximal at occipital electrodes around standard site Oz. (**b**) The difference in amplitude between the steady-state visual response to the left-and right-tilted gratings underwent clear changes in the direction of the Target type (higher-contrast in the left/right-tilted grating, thin/thick lines), and this directional effect varied in strength as a function of high/low Contrast difference (dark/light shades), and Speed/Accuracy Regime (red/blue). Specifically, there was an increased separation between the stronger sensory representation of the higher-contrast and the weaker representation of the lower-contrast grating under speed pressure. Upper panel shows the strength (F-value) of the Regime × Target type interaction in a time-resolved fashion, illustrating the time frame during which the differential between left-and right-targets was significantly widened under speed pressure (shaded grey). This differential effect of speed pressure was significant just before the response (F(1, 15)=5.7, p=0.031, Supplementary Table 2f) but ceased to be significant just after the response (F(1, 15)=2.9, p=0.11, Supplementary Table 2g). Note that the gradual initial increase/decrease in the differential SSVEP (−100 to about +200ms), despite the contrast change stepping instantaneously, is attributable to the fact that SSVEP amplitude at a given time is measured in a 400-ms window centered on that time, and this leads to the apparent deviation even before evidence onset (t=0).

Recent theoretical work has suggested that sensory-level modulations may be facilitated under speed pressure by increased cognitive effort^14,15^. We thus additionally examined measures of pupil size, which have been widely linked with effort and arousal^35,36^. We found relatively increased pupil size under speed pressure starting just before evidence onset, an effect which increased in magnitude over the course of evidence presentation (Figure 3). Moreover, greater pupil size predicted greater SSVEP differences between left and right targets even when the Regime effect is accounted for (Left/Right x Pupil: F(1,15)=6.07, p=0.026 Supplementary Table 2h).

**FIGURE 3:**
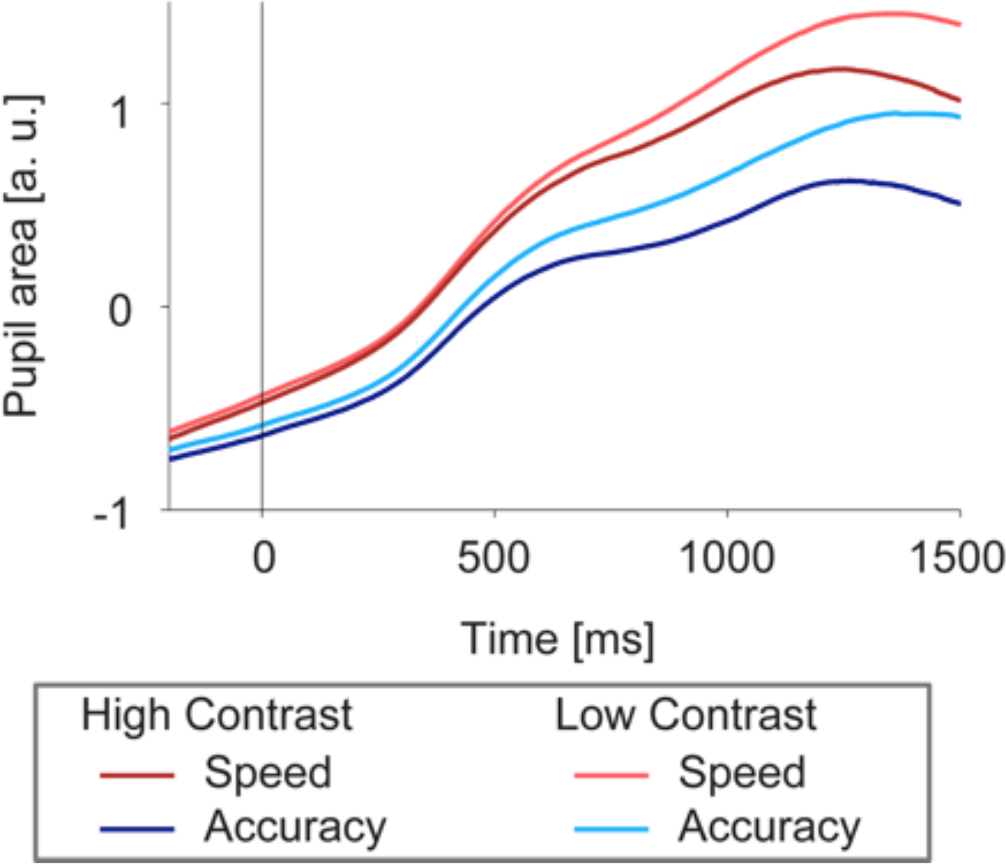
Pupil size is modulated by speed pressure. Traces depict mean pupil size plotted over time with respect to evidence onset. Traces are baseline-corrected to the 500ms before the onset of the regime-cue. Pupil size was increased under speed pressure (red traces) starting just before evidence onset (t(15)=2.25, p=0.040). This effect increased in magnitude over the course of evidence presentation, visible in the increasing separation between red and blue traces (two-way repeated-measures ANOVA, Speed/Accuracy by Time interaction, F(29,435)=17.6, p=0.00021, Supplementary Table 2j). A linear mixed-effects model of pupil size measured at response confirmed that pupil size was increased for Speed compared to Accuracy trials (t(14521)=14.01; p=2.5*10^−44^) while also accounting for an additional positive relationship with RT (t(14521)=9.92; p=3.8*10^−23^), and a negative relationship with Contrast (t(14521)=-7.80; p=6.54*10^−15^, Supplementary Table 1m).

### Urgency is applied directly to motor preparation but not to motor-independent accumulation

We traced the dynamics of decision-related buildup in Mu/beta-band indices of motor preparation over the motor cortex of each hemisphere, separately indexing each response alternative (Figure 4a-d)^23,30^, and also in a motor-independent signature of evidence accumulation (CPP; Figure 4e-h)^23,33^. As observed previously in continuous decision tasks that required immediate responses and hence the preparation of actions alongside evidence integration^23,33^, both signals exhibited a gradual, evidence-dependent build-up (Contrast effect on slope of Motor preparation for executed response: F(1,31)=9.2; p=0.008; CPP slope: F(1,31)=14.0; p=0.0026; see Figure 4c, 4g, Supplementary Table 2k, 2l) and reached their peaks around the time of the response. However, in this discrete-trial, urgent, forced-choice task there were several salient distinctions between the two levels.

Consistent with previous work^5–8,11^, motor preparation signals exhibited higher baseline activation prior to evidence onset following Speed cues (executed response: t(14969)=3.36; p=0.00079; withheld response: t(14969)=3.99; p=6.57*10^−5^ in linear mixed-effects models, Supplementary Table 1d-e). The trial-to-trial variability in this baseline or “starting point” of motor preparation was further predictive of RT, with lower levels predicting slower RTs (executed response: t(14969)=5.20; p=2.05*10^−7^; withheld response: t(14969)=2.59; p=0.0097; Supplementary Fig. 5A), which in the sequential sampling framework can explain the fast errors observed in the Speed regime^37^ (Figure 1c). Despite these variations in the baseline, and similar to sensorimotor neural activity in monkeys^38,39^, preparation towards the executed response (contralateral Mu/Beta) reached a stable threshold level just prior to response that did not vary significantly as a function of RT, Contrast or Speed/Accuracy regime (Figure 4c-d; linear mixed-effects model; all factors p>0.05, Supplementary Table 1f).

In contrast to motor preparation signals, the pre-evidence amplitude of the CPP, measured relative to a pre-cue baseline, showed no effect of speed pressure (linear mixed-effects model; p=0.22, Supplementary Table 1g) or relationship with RT (p=0.35). Further, CPP amplitude did not reach a stable level at the time of decision commitment but rather varied across conditions and reaction time (Figure 4h). Most strikingly, it decreased over RT in all conditions, aside from the very fastest, low-accuracy trials of the Speed regime, mirroring the most prominent feature of the conditional accuracy functions (Figure 1c). A linear mixed-effects model revealed that amplitude significantly decreased with reaction time (t(14696)=−3.4; p=0.00062, Supplementary Table 1h), increased with evidence strength (t(14696)=2.4; p=0.014), and showed a trend towards higher amplitudes in the Accuracy compared to Speed regime (t(14696)=1.7; p=0.09; Figure 4h). These results did not depend on the exact time window for measuring the amplitude at decision commitment (Supplementary Fig. 4) or whether a potential evoked by the auditory cue was removed (Supplementary Fig. 3).

**FIGURE 4:**
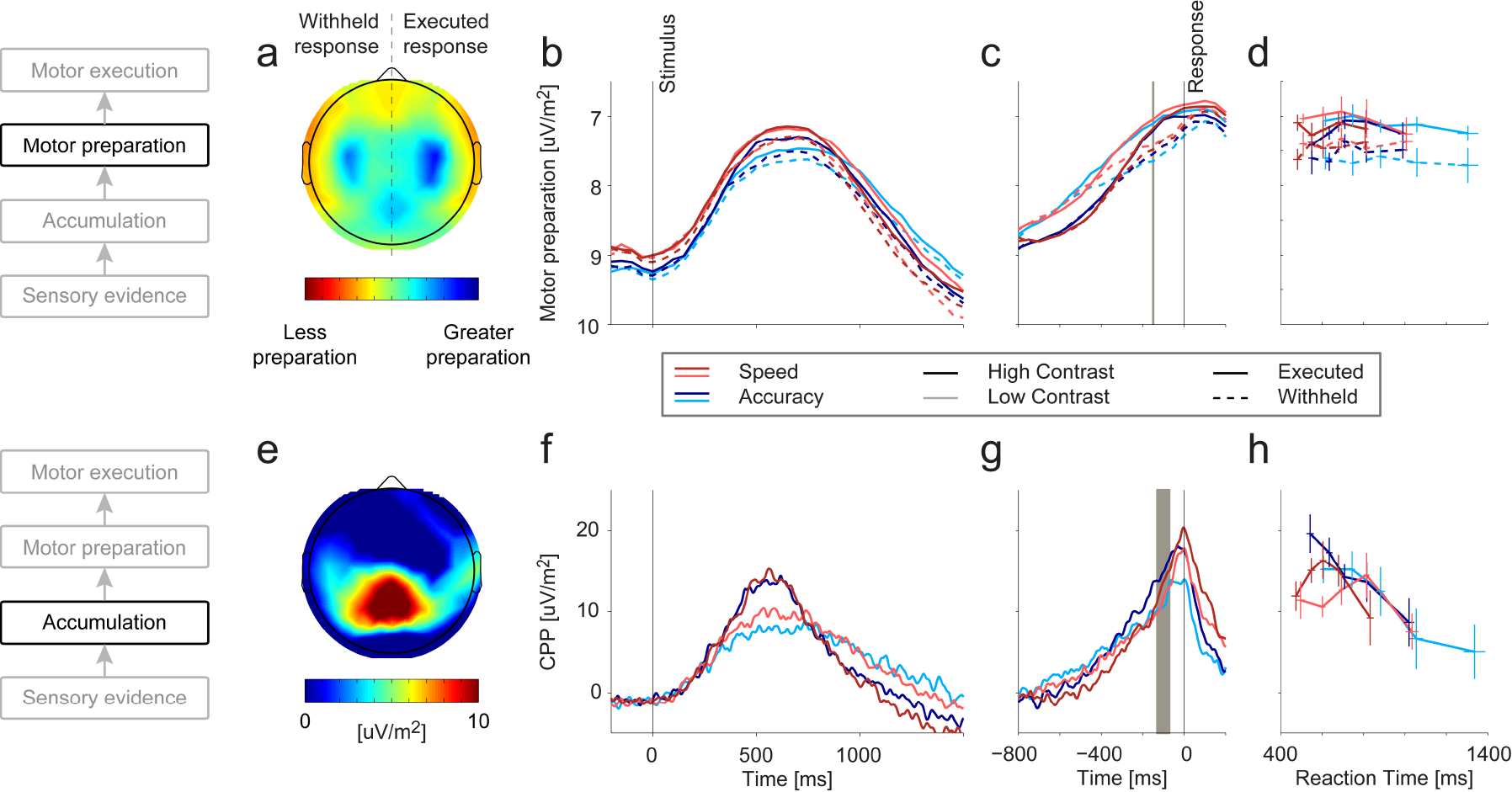
Effects of Evidence strength and Speed/Accuracy regime on effector-selective and motor-independent decision signals. **(a-d). Motor preparation. (a)** Topography of the decrease in Mu/Beta-band amplitude at response, relative to pre-evidence bafseline. Topographies for left-hand responses are collapsed with a left-right-reversed topography for right-hand responses, so that left/right scalp corresponds to motor preparation for the withheld/executed hand. **(b)** Stimulus-locked and **(c)** response-locked timecourses of Mu/Beta activity reflecting temporally increasing motor preparation (reduced spectral amplitude; note reversed y-axis) for the executed (solid) and the withheld (dashed) response. Spectral amplitudes for each timepoint are computed in a 300ms window centered on that timepoint, resulting in a temporal smoothing effect. The grey vertical line indicates the center of the 300-ms time window in which motor preparation at response was measured (from −300 to 0 ms). **(d)** Motor preparation at response plotted over response time. At response, the level of motor preparation for the executed response (solid) is independent of evidence strength (light vs. dark), Speed/Accuracy regime (red vs. blue), or reaction time. As expected, the motor preparation for the withheld response alternative ultimately reached significantly lower levels than that of the executed response (main effect of Executed/Withheld response, F(1,15)=14.2593; p=0.0018 Supplementary Table 2l). Error bars indicate S. E. M. **(e-h)**. **Motor-independent evidence accumulation. (e)** ERP topography around the time of response commitment (−130 to −70ms with respect to the button press, gray horizontal bar in g), showing a clear centro-parietal positivity. **(f)** Stimulus-locked and **(g)** response-locked centro-parietal traces for different evidence levels and Speed/Accuracy regimes, after subtraction of auditory evoked potential (see methods, Supplementary Fig. 3). **(h)** CPP amplitude around the time of decision commitment, plotted over response time. Note that although by the time of button click the CPP appeared to rise higher on average under the Speed than the Accuracy regime (G), this occurs clearly beyond any reasonable timeframe of decision commitment (Supplementary Fig. 4), and can be explained by a combination of a greater amount of post-decision accumulation under speed pressure (see EMG results section), and the greater impact of dynamic urgency at the later response times in the Accuracy Regime. Error bars indicate S. E. M. Note that y-axis scaling is identical across panels for each of the two decision signals, shown in panels B and F, respectively.

The lack of urgency effects on baseline CPP amplitude suggests that the motor-independent representation of cumulative evidence is spared from static urgency influences applied at the motor level. The decreased peri-response amplitude of the CPP for trials with longer RT further suggests that there is a dynamic component of urgency at the motor level, which is not applied directly at the CPP level. This follows from the fact that if, as the Mu/Beta signals suggest, the ultimate action-triggering threshold is set at the motor level, then increasing urgency at that level decreases the amount of evidence that can be accumulated before the motor threshold is crossed. Thus, motor-level dynamic urgency effectively “narrows the bounds” on the purely sensory-driven representation of cumulative evidence reflected in the CPP. Temporally increasing motor-level urgency is indeed reflected in Mu/Beta signatures of motor preparation in that both alternatives launch in the same direction towards threshold upon evidence onset (Figure 4b). Further, the “excursion” (level at RT relative to baseline) of preparation towards the unfavored alternative (ipsilateral Mu/Beta signal, dashed lines) increased over RT (t(13034) = −2.39; p=0.017), indicating that the longer a decision took, the more preparation was undertaken irrespective of cumulative evidence (linear mixed-effects model, Supplementary Fig.5, Supplementary Table 1i).

Based on the observed boost in differential evidence at the sensory level, a steeper build-up in both evidence accumulation and motor preparation within a trial would be predicted under Speed compared to Accuracy emphasis. Testing within-trial temporal slope measures prior to response, we indeed found a steeper buildup under Speed pressure both for the Mu/Beta motor preparation signals for the executed response (ANOVA on temporal slope; main effect of Speed/Accuracy regime F(1,15)=11.5; p=0.0040, Supplementary Table 2j) and for the CPP (F(1,15)=5.4; p=0.034, Supplementary Table 2k). Taken together, the pattern of effects on the CPP are consistent with two separate knock-on effects from adjustments made directly at other levels: the steepened buildup resulting from the boost at the sensory evidence level, and the curtailment of the CPP’s peri-response amplitude by motor-level urgency (see Figure 5).

**Figure 5:**
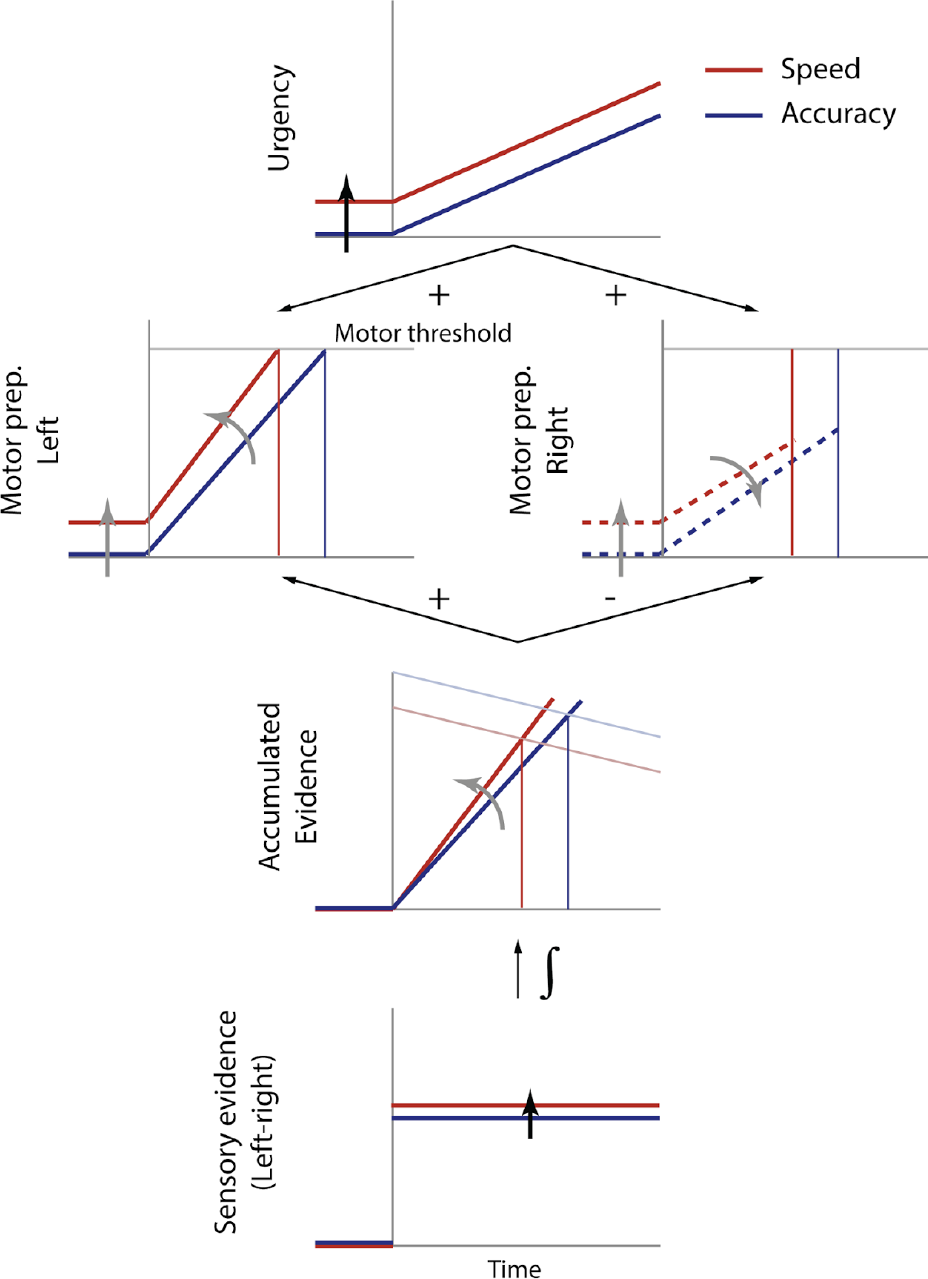
Schematic summary of speed pressure effects across cortical sensorimotor levels. In our proposed architecture, neural sensory evidence signals (bottom) are temporally integrated at an intermediate, motor-independent level (reflected in the CPP), which in turn feeds each of two competing motor-level buildup signals with opposite polarity for the alternative favored vs. unfavored by the cumulative evidence. An evidence-independent urgency signal (top) additively feeds both motor-level alternatives equally on average, and grows dynamically over time. The ultimate terminating and action-triggering threshold is set at this motor level, with the motor alternative that is first to reach its bound determining the response choice and timing. The schematic illustrates idealized dynamics at all levels for a typical single trial (left-tilted, correct) in each Regime (Speed in red, Accuracy in blue), representing the primary, active speed pressure adjustments by black arrows, and their knock-on impact at other levels by gray arrows. Based on the similar rates of decline in conditional accuracy (Figure 1c), we assume for parsimony that the rate of urgency buildup is the same for both Regimes, but the whole function is shifted upwards under Speed pressure (top). Meanwhile, at the sensory level (bottom), the differential representation of the increased versus decreased contrast grating is enhanced, constituting stronger sensory evidence. This does not constitute an urgency contribution according to its prevailing definition because it is evidence-selective, boosting the signal in the direction of the higher-contrast grating in the physical stimulus. These two primary adjustments have knock-on effects on the motor-independent level of evidence accumulation reflected by the CPP, without that level necessarily undergoing any direct adjustment itself: the enhanced sensory evidence leads to a slight steepening of buildup, while growing motor-level urgency results in an effective collapse in the attainable amplitude of the CPP by the time the motor-level threshold crossing triggers a response. Note that the scales of the axes at different levels are not intended to be consistent, and the size of some effects are exaggerated to aid clarity of illustration.

### Effects of Speed Pressure at the level of Muscle activation

We recorded electromyogram (EMG) signals bilaterally from the thenar eminence while subjects prepared thumb responses to report their decisions. Mean “motor times,” quantified as the time from the onset of the response-initiating EMG burst in the response-executing thumb to the button click, were significantly shorter under Speed compared to Accuracy emphasis (Figure 6a; t(15)=6.09, p=2.07*10^−5^). This was accompanied by a significantly steeper initial rise (linear mixed effects model; t(29398)=12.8; p=1.26*10^−37^, Supplementary Table 1j) and greater overall amplitude in muscle activation just prior (-100 to 0ms) to the executed response (Figure 6b-c; t(14696)=15.06; p=6.69*10^−51^, Supplementary Table 1k), in line with recent findings that the shortening of response-executing EMG bursts goes hand-in-hand with intensified muscle activation^40^.

The parallel signal recordings further allowed us to examine the dynamics of peri-response evidence accumulation within the muscle activation timeframe. In both regimes, the CPP peak time relative to button click completion (Figure 4g; Speed: −6.1 ± 39.6*ms*; Accuracy: −45.9 ± 37.6*ms*) was much later than EMG onset (Speed: −103.3 ± 16.7*ms*; Accuracy: −139.9 ± 17.6*ms*), suggesting that even for simple button presses, responses are initiated while the evidence accumulation process is ongoing^41,42^ (see Resulaj et al.^43^ for reaches). As might be expected given the accelerated muscle activation, the CPP peaked later relative to the click under Speed pressure than under Accuracy emphasis (F(1,15)=14.0; p=0.0019, Supplementary Table 2m). Interestingly, this delayed CPP peak under speed pressure was also observed with respect to EMG onset (2-Way ANOVA: F(1,15)=7.8; p=0.014, Supplementary Table 2n). Although the moment of decision commitment cannot be precisely ascertained, it can be assumed to occur no later than the peak response-executing EMG activation (−50ms), and the delayed peak of the CPP therefore suggests that there is significantly more post-commitment accumulation^44,45^ under Speed than Accuracy emphasis. Combined with the steeper buildup under speed pressure, this can explain why the CPP appears to rise to a higher peak following decisions under speed pressure in Figure 4g.

Consistent with recent reports^40,41^, significant muscle activation during decision formation was not confined to the effector ultimately producing the decision report; significant, “partial” (non response-completing) bursts of EMG activity could also be detected in the response-withholding thumb. Such EMG bursts were significantly more prevalent under Speed compared to Accuracy emphasis (Figure 6d; t(15)=4.6, p=0.00034). Further, EMG activation levels of the response-withholding thumb, measured immediately prior to the mean onset time of the response-executing burst to avoid spurious influences from the executing movement on the same mouse (−225 to −125ms relative to the button click), were significantly elevated under Speed emphasis (Figure 6e-f; t(14696)=5.5, p=3.66*10^−8^, Supplementary Table 1l) and increased over reaction time (Figure 6f; t(14696)=5.4, p=7.27*10^−8^). Since this is the hand to which the evidence is usually opposed, this latter result indicates that a dynamically growing urgency signal is manifest even at the peripheral stage of motor execution.

**Figure 6:**
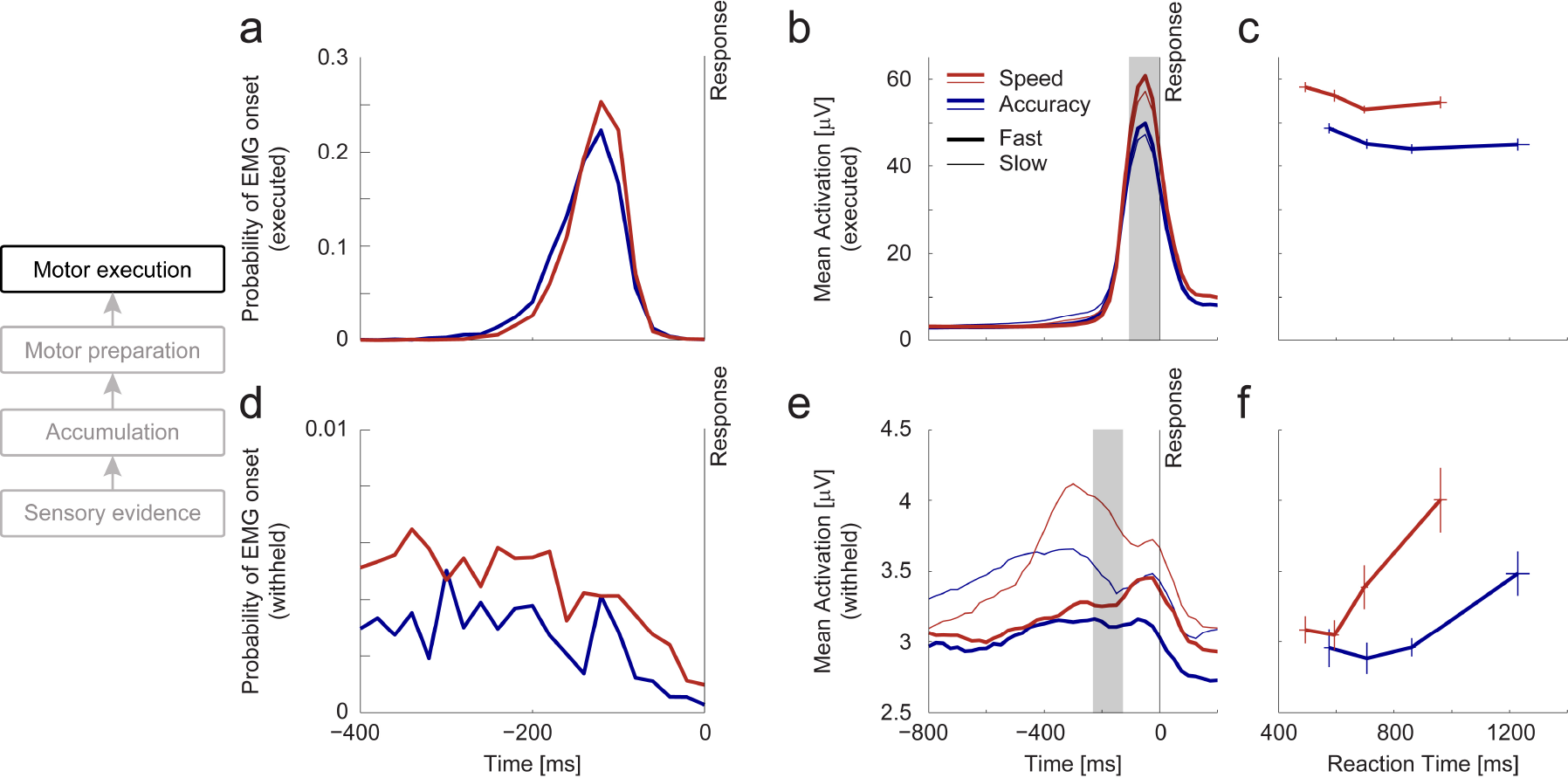
Urgency induces graded changes in peripheral muscle activation (a-c) Electromyographic (EMG) activity in the response-executing thumb. (**a**) Distributions of “motor times,” quantifying the time lag between muscle activation onset and action completion (button click), revealing shorter lags under Speed (red) than Accuracy (blue) emphasis. **(b)** Muscle activation (mean EMG spectral amplitude over 10-250Hz measured in 100ms-time windows) time-locked to the button click response is increased under speed pressure, and for fast (thick traces) compared to slower (thin traces) responses. **(c)** Muscle activation during response execution (100ms-time window before click, shaded gray in B) is increased under speed pressure across all response time bins, and decreases significantly over reaction time bins (Linear mixed-effects model: t(14696)=−5.6; p1.72*10^−8^, Supplementary Table 1k). Error bars indicate S.E.M. **(d-f) EMG activity in the response-withholding thumb. (d)** In the response-withholding thumb, the probability of a muscle activation onset occurring without triggering a motor response (“partial” burst) is increased under speed pressure across a broad time range. **(e)** Mean spectral amplitudes measured in two response time bins show that, especially for late responses (thin traces), response-locked traces of muscle activation in the response-withholding hand show increased activation under speed pressure (red traces). **(f)** With increasing response time, muscle activity in the response-withholding thumb increases both under Speed and Accuracy emphasis in a time window just prior to the mean EMG onset time in the response-executing thumb (−225ms to −125ms, shaded in E). Note that activation in the response-withholding hand is plotted on a much smaller scale than that of the response-executing hand. Error bars indicate S.E.M.

## Discussion

Our results reveal that speed pressure affects decision-related neural activity at each of the key processing levels necessary for contrast discrimination decisions, from the lowest cortical sensory level to the peripheral level of muscle activation. These modulations across the hierarchy arise from two principal adjustments that are fundamentally distinct in nature: an evidence-independent urgency contribution applied first at the motor preparation level which creates accuracy costs, and an enhancement of differential evidence at the sensory level that acts to alleviate those costs.

### Speed-accuracy adjustments at the sensory evidence level

Our finding of boosted differential sensory evidence under speed pressure stands in contrast to classical theoretical assumptions^2,4^, recent modeling results suggesting a lower quality evidence representation^16^, and fMRI studies finding no changes at the sensory level^7,19^. Given the transient nature of the sensory modulations we observed, it is possible that fMRI does not have the requisite temporal resolution to detect such effects, though differing task demands may also play a role. Our findings do broadly accord with the observation of earlier and stronger spatial selectivity for visual search targets under speed pressure in visual neurons of monkey FEF, although our results are distinct in a number of ways. First, while visual FEF neurons serve to represent the salience of items in their receptive field and thus furnish the evidence for visual search decisions^18^, our findings show that speed pressure can also impact on low-level representations of basic sensory attributes that form the evidence for simple discriminations requiring no spatial selection. Second, whereas speed pressure increased FEF activity somewhat indiscriminately before, during and at the end of decisions in both visual and motor neurons, our SSVEP modulation was strictly evidence-selective with no non-selective or baseline components to it (Supplementary Fig. 2). That is, the modulation served to widen the differential activity already driven by the bottom-up stimulus information, but not by “turning-up” the representation of both alternatives. This differential boost effect occurred alongside steepened accumulator signal buildup and in the absence of any apparent background noise modulation reflected in intervening frequencies (Supplementary Fig. 2), and it was linked with improved decision performance, all indicating that this modulation lessens the accuracy toll imposed by speed pressure rather than contributing to it.

There are several possible neural mechanisms that may underpin these evidence-selective changes. At the sensory level itself these include increased competitive interactions^46,47^, or an enhancement of the sensitivity of population contrast response functions in early visual cortex in the region of the 50% baseline contrast level, for example through the narrowing of orientation tuning centered on the two grating orientations^48^. Alternatively this modulation could come about through positive feedback from higher evidence-accumulation and/or motor levels, such that the representation of the currently favored alternative formed during early stages of accumulation is boosted^49^. Future work should aim to adjudicate between these possibilities.

The differential sensory evidence boost was accompanied by increased pupil size, and there was trial-to-trial covariation between the two. Pupil size has long been linked to generalized factors of effort and arousal^35,50^ and central neuromodulatory systems thought to support them, such as the Locus-Coeruleus Noradrenaline (LC-NA) system^13,51–54^ whose projection sites include sensory areas^55,56^. These neuromodulatory systems are thought to control global levels of neural gain^35^. Dynamic modulations in global gain during decision formation were suggested in theoretical work to play a role in optimizing decision making^25,57^, and recently the integration of empirical measurements of pupil size dynamics with computational models gave rise to the proposal that dynamic gain modulation may in fact be the generator of evidence-independent urgency^11^. Since gain modulation acts globally, an obvious prediction would be that even the sensory level is targeted by urgency influences. However, our observed sensory-level SSVEP modulations had no non-selective or evidence-independent component, wholly inconsistent with a core defining property of urgency. This does not preclude that the LC-NA system played a role in our sensory modulations; a growing number of theories of LC-NA function have asserted that its global influences may act to enhance selectivity, since the interaction of locally released glutamate and systemically released NA would act to enhance more active representations while suppressing less active ones^13,58–60^. Thus, our findings suggest that the impact of pupil-linked neuromodulatory systems on decision making may come in more forms than only accuracy-compromising urgency.

### A motor-independent representation of cumulative evidence spared from urgency

While both systematic and random, behavior-predictive variations were found in baseline amplitudes at the motor level, no such effects were observed at the motor-independent CPP level. Moreover, whereas an invariant action-triggering threshold was observed at the motor level, the CPP decreased in the amplitude it reached by the time of response for trials associated with greater levels of urgency, such as those with slower RT (Figure 4h). As illustrated in Figure 5, this can be explained as the knock-on effect of urgency influences operating at the motor level. In effect, the rising urgency signal at the motor level translates to a corresponding collapse in the attainable quantity of cumulative evidence at the CPP level. The effects on pre-response CPP amplitude were qualitatively mirrored in the conditional accuracy functions, consistent with the CPP reflecting an unadulterated representation of cumulative evidence. The quantitative trends in CPP and accuracy were by no means perfectly matched, however, which may relate to the form taken by the underlying evidence accumulation circuits that generate the CPP on the scalp. The CPP manifests as a positive deflection for either of the two decision alternatives in a motion discrimination task^33^ as well as in the current contrast discrimination task (Supplementary Table 1c), even when the incorrect response is chosen (t(15)=3.63, p=0.0024), and also for false target detections^23^, all indicating that the underlying neural evidence integration signals for *any* decision outcome contribute to the CPP in the same, positive direction^61^. This means that any proportional relationship between mean centro-parietal amplitude at response and response accuracy would break down when the latter approaches chance level. In particular, for longer-RT trials in the low-contrast condition, which would be characterized by weak evidence coupled with narrowed effective bounds, many errors may be associated with significant diffusion in favor of the incorrect alternative, which would translate to relatively elevated, positive CPP amplitudes at response even though response accuracy is greatly reduced.

An ongoing debate has centered on whether slow errors are, in general, better explained by a collapsing decision bound^62^ (equivalent to additive, evidence-independent urgency^8,9,11^) or by drift rate variability^2^ (e.g. see Hawkins et al.^63^ and Ratcliff et al.^3^). The collapsing CPP-amplitude effect observed here clearly points to the presence of the former mechanism. This does not preclude that there is some amount of drift rate variability, but renders it an unlikely primary driver behind slow errors. It is also noteworthy that this collapse in CPP amplitude contrasts strikingly with observed patterns in continuous monitoring tasks, where CPP amplitude is stable across RT in a similar way to motor signals^23,33^. In our proposed framework, this would indicate an absence of time-dependent dynamic urgency at the motor level in these tasks, which is plausible given that subjects are unable to predict the onset of the sensory evidence “targets” and, indeed, are unaware of missing them.

The decline in response accuracy (Figure 1c) and CPP amplitude at response (Figure 4h) occurred to a similar extent in the Accuracy and Speed regimes, suggesting that, despite an offset reflected in pre-evidence motor preparation levels, the rate of increase in urgency may be similar across the two regimes. This is likely due to the fact that the fixed stimulus duration imposed a response deadline even in the Accuracy regime, combined with the fact that difficulty levels were interleaved in both regimes, for which collapsing bounds represent the optimal strategy^12^. Moreover, in contrast with previous studies observing differences in urgency steepness across regimes^8,11^, here regime was randomly cued trial-by-trial rather than manipulated block-by-block.

Our finding of an invariant threshold level at response in effector-selective motor preparation signals accords with similar findings in monkeys^8,10^ and humans^11^ during feature discriminations, but this finding is not ubiquitous. For example, evidence accumulation signals reflected in FEF motor neurons during visual search decisions did not terminate at a stereotyped threshold level but rather exhibited increased termination levels under speed pressure^18^, running contrary to a central tenet of sequential sampling models and leading the authors to propose a further stage of integration beyond FEF. This might suggest that the task of visual search involves qualitatively different mechanisms for speed emphasis than simple discriminations.

Our demonstration that the brain computes a unitary representation of cumulative evidence that is spared from urgency influences at the motor level offers new insights into how choice-relevant information is represented in the brain. Abstract choice representations have been proposed as an efficient means for the brain to flexibly route sensory information to goal-relevant motor regions^20,21,33^ and, in fact, the suggestion was made in very early work that such signals may not be influenced directly by speed pressure^64,65^. We have previously demonstrated that, when speed pressure is absent and the onset of sensory evidence unpredictable, the evidence-dependent build-up of the CPP reliably precedes that of effector-selective signals^33^, suggesting that it intermediates between sensation and action preparation. Neurons in area LIP of the monkey have been found to encode goal-relevant stimulus categories but it is not known whether these signals exhibit evidence accumulation dynamics^21^. Moreover, while the CPP appears to rise solely as a function of cumulative evidence, LIP neurons multiplex a variety of task-and motor-related signals including urgency. Thus, the CPP appears to represent a decision variable signal of a different nature than the abstract signals identified to date in the monkey brain. Exactly how the two are related is a topic for future research.

### Urgency at the peripheral level of muscle activation

At the peripheral motor level, we found that muscle activation for even the unchosen response increased with elapsed time consistent with a dynamic urgency component of a similar nature to that found at the cortical level of motor preparation. Further, the time between the onset of substantial muscle activation and the button click was significantly decreased under speed pressure. At first glance, this appears consistent with a decrease in the additive “non-decision time” component of RT as found in computational model fits in some studies^66^. However a growing number of studies have demonstrated that action execution, even for simple button clicks, is not deferred until a decision bound is crossed but rather can be dynamically shaped by the ongoing evidence-accumulating decision variable^41,67,68^. The existence of partial EMG bursts in our task as well as others’^41^ underlines that EMG onset does not mark complete commitment or a “point of no return.” Assuming a fixed mapping of the decision variable to EMG activation, the decreased motor time could arise from either a decreased decision bound or steeper accumulation. The fact that our EMG signals rise more steeply and reach a higher amplitude under Speed pressure points to the latter explanation. This faster motor buildup may arise directly from the acceleration of the decision process under speed pressure due to enhanced sensory evidence encoding and potentially also from increased arousal. Whatever the mechanism, shortened motor time reduces the opportunity for retracting incorrect, partial responses^40,69^.

In contrast to our findings of shortened and intensified muscle activation, work in primates suggests no differences in saccade velocity as a function of RT or speed pressure^18^, implying that this finding does not necessarily generalize to all actions. While the muscle activations required to initiate saccadic responses stand in direct conflict with one another for different response alternatives, most other response modalities allow for the simultaneous preparation of multiple responses at the muscular level with a much lower degree of antagonism^70^. The presence of significant muscle activation and discrete EMG bursts in the response-withholding effector in our data presents strong evidence that subjects were indeed preparing both responses simultaneously, and to a greater degree under speed pressure.

Taken together, our results serve to highlight the value of the global systems-level view over decision making mechanisms afforded by noninvasive assays in humans employed in the context of purpose-designed paradigms. We uncovered multifaceted adaptations to speed pressure at the sensory, evidence accumulation, motor preparation and motor execution levels as well as dynamic neuromodulatory influences reflected in pupillometry. As demonstrated here, multi-level signal tracking affords the ability to test predictions of a hypothesis at multiple hierarchical processing levels, or to adjudicate between alternative interpretations of an effect at one level by testing predictions that apply to other levels. More generally, our findings underscore the emerging imperative to move from one-dimensional decision models to more neurally-based models embracing the hierarchical, interactive, and flexible nature of real neural systems accomplishing adaptive decisions^1,24–27^, and highlight that neural recordings in humans can act as a strong guide.

## Online Methods

### Participants

Sixteen participants (five male, *mean* ± *std* = 27.06 ± 4.72) gave written and informed consent to partake in this study. All participants had normal or corrected-to-normal vision, and no history of psychiatric illness or head injury. All procedures were approved by the Institutional Review Board of the City College of New York and were in accordance with the Declaration of Helsinki. Subjects gave informed consent and were compensated for their participation with between $9/hour and $15/hour depending on their task performance.

### Contrast discrimination task

Participants were asked to perform a discrete two-alternative forced-choice contrast discrimination task. Visual stimuli were presented on a gamma-corrected CRT monitor (Dell M782) with a refresh rate of 100 Hz inside a dark, sound attenuated, and radio frequency interference-shielded room. Visual and auditory stimulus presentation was programmed in Matlab (MATLAB 6.1, The MathWorks Inc., Natick, MA, 2000) using the PsychToolbox extension^71,72^. Participants were seated at a viewing distance of 57 cm from the monitor, and asked to fixate on a central fixation point. The participants’ task was to judge which of two overlaid, orthogonal gratings was greater in contrast. The imperative stimulus was an annular pattern with an inner and outer radius of 1° and 6° of visual angle, respectively, presented centrally on a gray background with the same mean luminance (65.2 cd/m^2^). The pattern consisted of two overlaid square-wave gratings (spatial frequency = 1 cycle per degree) tilted at −45 and +45 degrees relative to vertical, which were phase-reversed at 20Hz and 25Hz, respectively (Figure 1a). After 200ms of presenting just the fixation point (2x2 pixels), trials began with a linear fade-in of this stimulus from 0% contrast to 50% contrast of both gratings over 800ms at the end of which a reward-regime cue was presented in the form of a change in the color of the fixation point (colors randomized across subjects). This regime cue remained on the screen throughout the duration of the trial. During the following baseline stimulus presentation (800, 1200 or 1600 ms in a pseudorandomized order) the contrast of both gratings was held at 50%. The contrast of one grating was then stepped up to 56% (low) or 62% (high) and that of the other grating simultaneously stepped down to 44% (low) or 38% (high), and this contrast differential was held fixed for a full 2400-ms evidence-presentation interval. Evidence onset was marked by a simultaneously onsetting 100 ms-tone (10 kHz, 5ms fade-in/fade-out) to exclude any temporal ambiguity about evidence onset. Participants indicated that the left-tilted or right-tilted grating was higher in contrast by clicking a mouse button with the thumb of the corresponding hand. At the end of this interval, feedback was provided in the form of the number of points won on the current trial presented close to fixation for 200ms alongside a tone whose pitch was proportional to this number of points and whose length indicated whether the response given had been correct (100ms/250ms beep for correct/incorrect responses, double beep for responses after deadline). After every 10 trials, participants received an extended feedback in the form of an information screen stating the number of points won on the last 10 trials as well as the number of points won in the current experimental block to that point. Within experimental blocks each trial’s evidence-onset delay, target-direction, contrast level, and Speed/Accuracy regime (see next paragraph) was assigned pseudorandomly.

### Speed/Accuracy regimes

This contrast discrimination task was performed in four different reward regimes, where particularly fast or accurate responses were rewarded highly on Speed or Accuracy-trials, respectively. Reward conditions included 1) response time (RT)-independent rewards up to a late deadline which coincided with evidence offset (‘Accuracy deadline’), 2) RT-independent rewards up to an early deadline at 1s after evidence-onset (‘Speed deadline’), 3) RT-dependent rewards, which decreased at a low rate (−4.2pts/s, ‘Accuracy slope’), and 4) RT-dependent rewards, which decreased at a high rate (−50pts/s, ‘Speed slope’). Rewards as a function of response time are displayed in Supplementary Fig. 1A-B. These reward regimes were initially designed to enable exploration of differences in speed-adaptation mechanisms in the decreasing-reward versus deadline regimes. Through extensive piloting the exact temporal deadlines and rates of reward decrease were adjusted to match mean reaction times across the two different methods of Speed/Accuracy emphasis manipulation. In each experimental block, the two regimes for a single reward manipulation method were randomly interleaved for comparison (i.e., either ‘Accuracy deadline’ vs. ‘Speed deadline’, or ‘Accuracy slope’ vs. ‘Speed slope’). 15 participants completed 16 blocks of 60 trials, and one subject completed 24 blocks of 40 trials. Subjects were instructed to try to maximize their points won in every experimental block, as their monetary reward after the experiment was calculated as a function of the sum of points won in four randomly chosen blocks.

### Behavioral analysis

Participants’ behavior was evaluated based on reaction time (RT) distributions for correct and incorrect responses. As an initial step, we determined whether there was a significant difference in reaction time distributions between the deadline and slope conditions of the two different Speed/Accuracy regimes (‘Speed deadline’ vs. ‘Speed slope’, and ‘Accuracy deadline’ vs. ‘Accuracy slope’) using Kolmogorov-Smirnov tests. Since these tests revealed no significant differences between RT distributions in any experimental condition, all consecutive analyses were performed on data pooled across the deadline and slope methods, but separately for Speed and Accuracy emphasis. All patterns of results were, however, verified for the individual deadline and slope methods. Conditional accuracy functions were computed as the proportion of correct trials within reaction time deciles. To examine potential differences in the rate of decline of accuracy over the slower-RT trials for which this decline was evident, we identified trials with RTs slower than the mean RT in the RT-bin with the greatest performance in each individual’s conditional accuracy function (trials pooled across conditions). We then performed a linear mixed-effects analysis on these trials, with fixed effects of RT, Speed/Accuracy emphasis, and Contrast, and Subject identity as a random effect. A likelihood ratio test was performed to determine whether the inclusion of an interaction between RT and Speed/Accuracy emphasis significantly improved model fits to the data.

### Data acquisition and pre-processing

Continuous Electroencephalogram (EEG) and Electromyogram (EMG) were acquired using a 96-channel actiCAP system and Brain Products DC amplifiers (Brain Products GmbH, München, Germany) at a sample rate of 500 Hz. 93 channels were used for a customized EEG montage including standard site FCz used as the online reference. The remaining four electrodes were used for recording EMG from the thenar eminence of the left and right thumb. Simultaneously, eye gaze and pupil size were acquired continuously using an EyeLink 1000 (SR-Research) eye tracker. Data were analyzed offline using in-house Matlab scripts in conjunction with data reading routines and topographic mapping functions of the EEGLAB toolbox^73^.

EEG and EMG data were detrended linearly offline within each experimental block. Potentials in each EEG electrode were further re-referenced to the average of all EEG channels, a Hamming low-pass filter with a cutoff frequency of 45 Hz was applied, and noisy channels were detected based on their elevated signal variance with respect to the rest of the channels, and interpolated for individual blocks using spherical spline interpolation (an average of 0.67 ± 0.46 channels per block). Individual trials were rejected from the analysis if the amplitude of a channel of interest exceeded 90 V or any electrode’s potential exceeded 180 V at any time point before the response. All EEG data were converted to current source density (CSD)^74^ to increase spatial resolution and specifically to reduce spatial overlap between the centro-parietal positivity and the fronto-central negativity^33^. Event-related potentials were then extracted from the EEG, EMG and pupillometry data for two different epochs: regime cue-locked epochs were extracted from 500 ms before the onset of the regime cue to 4000 ms thereafter, and target epochs spanned the 1000 ms before and the 3200 ms after evidence onset. EEG epochs were baseline-corrected relative to the 100 ms interval preceding the regime-cue, or evidence onset, respectively, and trials were rejected if the delay between the visual contrast change on the screen and the tone marking this evidence onset for the participant exceeded 30 ms, which occurred on less than 2.5% of trials. Due to the longer time constants associated with changes in pupil size, event-related pupil size waveforms were baseline-corrected with reference to the 500 ms prior to the onset of the regime-cue for both the cue-locked and the evidence onset-locked epochs. Response-aligned traces in all modalities were derived by extracting epochs from -1000 ms to 600 ms relative to the response on every trial. Spectral EEG measures were extracted from both stimulus-aligned epochs and response-aligned epochs through short-term Fourier transforms using boxcar windows of 300 ms (8 to 30Hz) or 400 ms (20 and 25 Hz) measured in steps of 50 ms.

### Signal Analysis

#### Sensory evidence representation

The cortical representation of visual evidence was quantified as the difference in Steady-State Visually Evoked Potentials (SSVEP) of the two target frequencies of 20 and 25 Hz averaged together with their respective first harmonics, and normalized to their respective neighboring frequency bins at standard site ‘Oz’. SSVEPs were measured on a single-trial basis using a standard short-time Fourier transform with a boxcar window size of 400ms, fitting an integer number of cycles of both the 20-Hz and 25-Hz components, with a step size of 50ms. 3-Way repeated-measures Analyses of Variance (ANOVAs) were used to determine the effect of the imposed Speed/Accuracy regime, stimulus Contrast (high vs. low) and Target Type (left-tilted increase vs. right-tilted increase) on the differential evidence signal (20 Hz minus 25 Hz spectral amplitude) in a time range of 250-450 ms post evidence onset. Sphericity was confirmed for all inputs to this and all other ANOVAs using Mauchly’s test. For visualization, these SSVEP amplitudes were baseline-corrected to the 400 ms preceding evidence onset. Having established a differential modulation effect in a Regime x Target Type interaction in this timerange, we examined the temporal extent of the effect by repeating the same ANOVA on all individual time windows after evidence-onset, and plotting the timecourse of F-values (Figure 2b, top). To determine whether the observed SSVEP modulations effects were specifically invoked during decision formation related to active stimulus evaluation, we repeated the same ANOVA on a response-locked time windows just prior (−50ms) and just after the button click (+50ms). On raw SSVEP amplitudes of each individual SSVEP frequency we carried out ANOVAs with the same factors, so that common, or non-selective effects could be assessed through main effects of Speed/Accuracy Regime (Supplementary Fig. 2).

#### Evidence accumulation

A centro-parietal positivity (CPP) previously linked to evidence accumulation (O’Connell et al 2012) was measured at standard site ‘Pz’. Because the onset of evidence was marked by a readily-audible tone, which itself generates a stereotyped auditory evoked potential, we employed an iterative algorithm for signal decomposition to separate out any strictly stimulus-locked auditory component from the decision-related signals based on the variability in response latency across single trials^75^. Critically, the algorithm was naive to all stimulus conditions, and the resulting stimulus-locked component was constrained to be invariant across all trials within each individual subject, so that differences in potentials evoked by stimulus contrast or Speed/Accuracy regime were left untouched by this method. This stimulus-locked component was then subtracted from the evoked potential of all trials.

From the resulting **centro-parietal positivities** we obtained three single-trial measures. The level of baseline activity was quantified as the average potential in the 50 ms before evidence onset (measured in the regime cue-locked epochs), and statistically assessed via a linear mixed-effects model. This as well as all following mixed-effects models included the fixed effects factors of Speed/Accuracy regime, Contrast difference, Reaction Time, squared Reaction Time, and Response hand or Target type (left vs. right), and Subject identity was always included as a random effect. We adopted the blanket policy of modeling any potential differences between left and right target trials because the two gratings had different phase-reversal frequencies, and similarly always included the RT-squared term based on the observation that conditional accuracy functions followed an inverted-U shape, suggesting potentially separate mechanisms at work to explain the lower accuracy in the fastest, versus the slowest RT trials. Neural measures and reaction times were always z-scored within subjects before being entered into the models. The rate of rise of the CPP was measured as the slope of the response-locked traces between -300 and -50 ms with respect to the response, chosen to capture the period of evidence accumulation on the vast majority of trials. The impact of stimulus Contrast and Speed/Accuracy regime on this rate of rise was established through a 2Way repeated measures ANOVA. Statistical test outcomes did not depend on the exact choice of this time window. The CPP amplitude at response was measured in a 60-ms window around the inflection point of the lateralized motor potential traced over contralateral motor cortex (-130 to -70ms, Supplementary Fig. 4), and statistically assessed via a linear mixed-effects model. The exact time window used for the CPP amplitude analysis did not influence the results qualitatively. These analyses were repeated using the raw-EEG data without subtracting the auditory evoked component, and results remained qualitatively unchanged (see Supplementary Fig. 3). To assess whether the CPP amplitude at response was significantly greater than zero when subjects made incorrect decisions, we pooled error trials across experimental conditions and computed a repeated-measures ANOVA across subjects. The peak time of the mean CPP within each subject, evidence-level and Speed/Accuracy regime was measured as the maximum amplitude between −150ms and +100ms relative to the response of a smoothed average trace. Smooth traces were computed by applying local regression using weighted linear least squares and a first degree polynomial model to moving windows of 200ms. A 2-Way ANOVA was computed to determine the effect of Speed/Accuracy regime and evidence-strength on this delay.

#### Motor preparation

Motor preparation signals were measured in the decrease of Mu/Beta amplitude (8-30Hz; integrated across both bands as in^32,34^) at motor cortical sites ‘C3’ (left) and ‘C4’ (right) for the preparation of contralateral responses. Spectral amplitude was quantified using a standard short-time Fourier transform with a boxcar window size of 300ms at intervals of 50ms. Motor preparation at baseline activation and at response were quantified as the Mu/Beta amplitude in the 300 ms preceding evidence onset and the button click, respectively, separately for the hemisphere contralateral and ipsilateral to the eventually executed response on a single trial basis. All measures were statistically assessed via linear mixed-effects models. To test for signatures of evidence-independent components of motor preparation that grow over time, we computed a linear mixed-effects model on the signal excursion of Mu/Beta amplitude ipsilateral to correct responses, where “excursion” is computed as the difference between levels at response and at the pre-evidence baseline. It was important to measure excursion in this case in order to account for the significant variation of baseline motor preparation with RT, which otherwise could obscure the influence of systematic urgency increases during evidence accumulation. Excluding the SSVEP frequencies (20 Hz and 25 Hz) from the Mu/Beta computations did not change the pattern of results. The rate of change in motor preparation was calculated on a single trial basis by measuring the slope of a line fit to the Mu/Beta amplitude in the interval between −350 and −150 ms relative to the response, chosen to capture as long a section of decision formation as is feasible while avoiding influences of post-response changes due to temporal blur associated with the 300-ms windows. A repeated measures ANOVA was computed to test the significance of the influence of stimulus Contrast level and Speed/Accuracy emphasis on the Mu/Beta slope contralateral to response for trials that resulted in a correct response only. Restricting this analysis to correct trials ensured a positive relationship between the physical evidence and the neural measure of motor preparation.

#### Response execution

EMG data were analyzed for effects on movement onset times and muscle activation levels. Motor time was quantified on a single-trial basis as the time between the onset of the muscle activity burst closest to response and the registration of a button click. EMG onset bursts were identified manually by visual inspection of the raw data, using a custom-made Graphical User Interface, and the results were verified on times derived from an automated algorithm relying on changes in variance of the broadband EMG signal and on the times estimated by another, independent human observer. The difference in mean motor time between Speed/Accuracy regimes was assessed using a two-tailed t-test. The effect of stimulus Contrast and Speed/Accuracy regime on the mean delay between EMG burst onset and the peak of the CPP were evaluated by 2-Way repeated measures ANOVAs. Muscle activation was quantified as the mean spectral amplitude between 10 and 250 Hz in 100-ms time windows stepped by 25ms. The ultimate response-producing muscle activation was quantified as the spectral EMG amplitude in the responding thumb in the 100ms preceding the button click. Insight into evidence-independent components of muscle activation was sought by quantifying the mean EMG spectral amplitude in the response-withholding thumb in a 100-ms time window preceding the mean onset time of the response-producing EMG burst (−225ms to −125ms). In this latter analysis only correct trials were included so that the sensory evidence runs counter to the measured action alternative, enabling us to more confidently attribute any increase over RT to evidence-independent urgency. Both measures were evaluated by linear mixed effect models.

On a single-trial basis, we additionally measured the rate of building muscle activation during responses initiation. Specifically, we measured the slope of a line fit to the spectral muscle activation timecourse (as before but stepped by 5 ms for increased resolution) in the response-executing thumb in the interval between −175ms and −125ms relative to the button click. This interval was chosen based on visual inspection of grand-average traces in single subjects. We statistically tested these temporal EMG slope measures using linear mixed effect models.

#### Pupillometry

To examine the role of pupil-linked arousal systems in the speed pressure adaptations, we continuously measured pupil size using the eye tracker. Pupil size was compared across Speed/Accuracy regimes in the pre-evidence baseline by a t-test. To test the influence of time on pupil size, we computed mean pupil size in 30 50-ms time windows spaced at 50ms starting at stimulus onset. We then tested these time series for a significant interaction between Speed/Accuracy emphasis and Time across subjects using a two-way (2x30), repeated measures ANOVA. We further measured pupil size at response in a 100-ms time window centered on the time of the button click and assessed it statistically via a linear mixed-effects model. In order to test whether, above and beyond the average adjustments for speed pressure, variations in pupil size were linked with variations in the differential evidence representation, we split the trials in each individual condition into two pupil-size bins based on mean pupil size between 0 and 1500ms after evidence onset. We then computed a 4-Way ANOVA including factors of Speed/Accuracy Regime, Contrast, Target type (Left/Right), and Pupil size to capture the effect of pupil size on the differential SSVEP in a Pupil × Target type interaction. Here differential SSVEP was measured in the same time frame during which the Speed/Accuracy effect was significant (200-550ms).

## Acknowledgements

This study was supported by grants from the United States National Science Foundation (BCS-1358955 to S.P.K. and R.G.O.), the European Research Council (63829 to R.G.O.) and the European Recovery Program-fellowship by the German National Merit Foundation (to N.A.S.). The authors are grateful for substantial feedback they received on the manuscript from Isabel Vanegas, Peter Murphy, Ariel Zylberberg, Dave McGovern and Ger Loughnane.

## Author contributions

NS, RGO and SPK designed the experiments. NS conducted the experiments and analyzed the data. NS, RGO and SPK wrote the paper.

## Competing financial interests

The authors declare no competing financial interests.

